# BDNF augmentation reverses cranial radiation therapy-induced cognitive decline and neurodegenerative consequences

**DOI:** 10.1101/2024.09.23.614590

**Authors:** Sanad M. El-Khatib, Arya R. Vagadia, Anh C. D. Le, Ding Quan Ng, Janet E. Baulch, Mingyu Du, Zhiqun Tan, Xiangmin Xu, Alexandre Chan, Munjal M. Acharya

## Abstract

Cranial radiation therapy (RT) for brain cancers is often associated with the development of radiation-induced cognitive dysfunction (RICD). RICD significantly impacts the quality of life for cancer survivors, highlighting an unmet medical need. Previous human studies revealed a marked reduction in plasma brain-derived neurotrophic factor (BDNF) post-chronic chemotherapy, linking this decline to a substantial cognitive dysfunction among cancer survivors. Moreover, riluzole (RZ)-mediated increased BDNF *in vivo* in the chemotherapy-exposed mice reversed cognitive decline. RZ is an FDA-approved medication for ALS known to increase BDNF *in vivo*. In an effort to mitigate the detrimental effects of RT-induced BDNF decline in RICD, we tested the efficacy of RZ in a cranially irradiated (9 Gy) adult mouse model. Notably, RT-exposed mice exhibited significantly reduced hippocampal BDNF, accompanied by increased neuroinflammation, loss of neuronal plasticity-related immediate early gene product, cFos, and synaptic density. Spatial transcriptomic profiling comparing the RT+Veh with the RT+RZ group showed gene expression signatures of neuroprotection of hippocampal excitatory neurons post-RZ. RT-exposed mice performed poorly on learning and memory, and memory consolidation tasks. However, irradiated mice receiving RZ (13 mg/kg, drinking water) for 6-7 weeks showed a significant improvement in cognitive function compared to RT-exposed mice receiving vehicle. Dual-immunofluorescence staining, spatial transcriptomics, and biochemical assessment of RZ-treated irradiated brains demonstrated preservation of synaptic integrity and neuronal plasticity but not neurogenesis and reduced neuroinflammation concurrent with elevated BDNF levels and transcripts compared to vehicle-treated irradiated brains. In summary, oral administration of RZ represents a viable and translationally feasible neuroprotective approach against RICD.

## INTRODUCTION

With the growing annual incidence of cancer diagnoses, an increasing number of patients are undergoing conventional cancer treatment modalities including radiation therapy (RT). Whole and partial brain RT is a primary treatment modality for most brain cancers to improve the overall survival, palliation or to control for the metastatic progression. In the United States, approximately 100,000 brain cancer patients receive RT every year and about 50 to 90% of survivors display cognitive dysfunction [15, 48]. While RT has demonstrated efficacy as a therapeutic intervention, its unintended consequences, including memory loss, impaired concentration, cognitive processing difficulties, slow response speed, and compromised executive functioning, persist among patients post-RT [24, 34, 45]. Clinical reports indicate that these long-term side effects result from enduring structural changes in the brain post-RT [34]. The presence of prolonged and debilitating consequences of RT in the medical landscape underscores the need for developing translationally feasible therapeutic approaches to mitigate cognitive decline and improve patients’ overall quality of life [38].

Clinical investigations have revealed a noteworthy correlation between the levels of brain-derived neurotrophic factor (BDNF) and cognitive outcomes in cancer patients exposed to systemic chemotherapy [39, 40]. A breadth of literature has established the link between lower BDNF levels in individuals diagnosed with neurodegenerative disorders such as Huntington’s disease (HD), Alzheimer’s disease (AD), and multiple sclerosis (MS) [10]. Synthesized in both neurons and glia, BDNF plays a multifaceted role, encompassing functions such as the maturation and development of neurons, axonal and neurite outgrowth, neuronal repair, neurotransmitter release or uptake, synapse strengthening, facilitation of long-term potentiation (LTP), regulation of plasticity, and interaction with the inflammation signaling [10, 12, 21]. Neuronal plasticity, a crucial aspect of adaptive neural functioning, enables neurons to respond to new environmental conditions through both functional and structural modifications. However, the disruption of these neuronal functions during conventional whole-brain RT was linked with cognitive dysfunction [7, 32] and demonstrates the importance of BDNF in its pathogenesis. Based on the previous scientific premise, we hypothesize that RT-induced impairments in neuronal function, elevated neuroinflammation, and cognitive deficits culminate following reductions in the neuroprotective BDNF.

The primary rationale for improving *in vivo* BDNF levels in patients with neurodegenerative conditions such as AD, HD, and MS is to safeguard the brain against substantial structural alterations when exposed to the non-conducive, neurodegenerative environment associated with the disease. In this context, an *in vivo* increase in BDNF following oral administration of riluzole (RZ) improved cognitive function in a mouse AD model [19]. A comparable approach can be applicable to cancer-related cognitive impairments. Our prior investigations have demonstrated that the oral administration of RZ via drinking water enhanced brain BDNF levels in an animal model subjected to cytotoxic chemotherapy [43]. RZ-mediated increases in BDNF ameliorated chronic chemotherapy (doxorubicin)-induced decline in cognitive function, neurogenesis and prevented neuroinflammation. RZ is an orally bioavailable, and FDA-approved treatment for ALS [47]. In this study, we posit that the administration of RZ will similarly elevate BDNF levels and transcripts, hence protecting the brain from cranial RT-induced synaptic loss and neuroinflammation and improve cognitive function.

## MATERIALS AND METHODS

Comprehensive experimental methods, materials, cognitive function, and tissue analysis protocols are provided in the **Supplemental Information** section.

### Animals and RZ treatment

All animal experimentation procedures were ethically approved by the Institutional Animal Care and Use Committee (IACUC) and conducted in compliance with the guidelines established by the National Institutes of Health (NIH). 12-13 weeks old male wild-type mice (C57BL/6) were procured from Jackson Laboratories and were group-housed (four mice per cage) under standard conditions, including a 12-hour light-dark cycle, room temperature maintained at 20 °C ± 1, and humidity at 70% ± 10. The mice were provided standard rodent chow diet (Envigo Teklad 2020X) by the University Laboratory Animal Resources (ULAR) at the University of California, Irvine. Previous studies suggested that adult female mice are resistant to acute (9-10 Gy) cranial RT-induced cognitive decline and neuroinflammation [17, 18]. As a proof of principle to support a neurotrophic factor’s influence on the damaged irradiated brain, for this study we focused on male mice. During the first week, mice were anesthetized (5% induction and 2% maintenance isoflurane gas) and cranially irradiated (9 Gy or 0 Gy sham irradiated) using Small Animal Radiation Therapy (SmART) equipment (Precision, Inc.). Mice were divided into the following groups (N=12-24 mice/group): Control or irradiated receiving vehicle (Con + Vehicle, RT + Vehicle), or RZ (Con + RZ, and RT + RZ). RZ (2-amino-6-(trifluoromethoxy) benzothiazole, Selleckchem) was dissolved in filter-sterilized (0.2 µm, Millipore), warm reverse osmosis (RO) water with constant stirring (1 to 2 hours at 45 °C) to achieve a stock concentration of 600 µg/ml [19]. The stock solution was stored frozen at -20 °C until usage. The working solution (60 µg/ml) of RZ was prepared twice a week by diluting the stock with filter-sterile RO water. Throughout the study, mice had ad libitum access to either the vehicle (filter-sterile RO water) or Riluzole solution (13 mg/kg per mouse, per day, as previously described) [19, 43]. To assess the impact of RZ treatment on *in vivo* dentate neurogenesis, two weeks post-RT, mice were administered BrdU (5-bromo-2’-deoxyuridine, 50 mg/kg, IP, Sigma) in phosphate-buffered saline (PBS, 100 mM, pH 7.6, Sigma) once daily for six days.

### Cognitive function testing

To investigate the impact of RZ on cognitive function post-RT, mice underwent cognitive assessments and anxiety-related behavioral tests four weeks following RZ treatment. Detailed test protocols are available in the Supplemental Information section. The cognitive function tasks, spanning two to three weeks, encompassed the open field test (OFT), and elevated plus maze (EPM) for anxiety-like behavior, object location memory (OLM), and a fear extinction memory consolidation task (FE). The OFT task determined the exploration of animals in an open arena to compare central (60% of central arena) and peripheral zones of the arena. The EPM test assesses an animal’s exploration of elevated open arms versus closed (dark) arms in a brightly lit environment. Anxious animals were expected to spend more time in closed arms than open arms. The hippocampal-dependent OLM task evaluated episodic and spatial memory function (9-10), and gauged an animal’s capacity to explore novel object placements in an unconfined, non-invasive open environment (arena) with bedding. The performance on the OLM task is quantified as the Memory Index (MI = [Novel/Total exploration time] – [Familiar/Total exploration time]) × 100. A positive MI denoted a preference for exploring novel spatial locations, while a negative or zero index indicated little or no preference, with equal or less exploration times for familiar and novel places. Following the completion of the OLM, with an intermission of approximately 72 hours, animals were subjected to the FE task. This task aimed to discern the influence of cranial RT or RZ treatment on hippocampal-dependent fear conditioning and the subsequent memory consolidation process (extinction memory), which actively involves dissociating learned responses to past adverse events. Briefly, during the conditioning phase, mice encountered three pairs of evenly spaced auditory stimuli coinciding with a mild foot shock. Twenty-four hours later, across the subsequent three days (extinction training phase), animals were exposed to 20 tones within the same contextual environment, including the odor and visual cues. 24 hours after completion of extinction training, the fear test was administered, during which animals encountered three tones in the same context. The freezing behavior of the animals was captured using a ceiling-mounted camera in the FE test chamber and analyzed using an automated freezing measurement module (FreezeFrame, Coulbourn Instruments). The percentage of time each mouse spent freezing during the tone was then calculated for the conditioning, extinction training, and testing phases. For the extinction training phase, an average five tones and four data points per day are presented. Higher the freezing on FE task indicates compromised fear memory consolidation process. The integration of the data from these tests equips us with robust analytical tools for assessing the impact of cranial RT and RZ on cognitive function. Further details of the protocols are provided in the **Supplemental Information** section.

### Immunofluorescence staining, confocal microscopy and immunoreactivity quantification

Following the completion of cognitive function tests, mice were euthanized via intracardiac perfusion using saline with heparin (10U/ml, Sigma) and 4% paraformaldehyde (PFA) prepared in 100 mM PBS, pH 7.4 (Sigma). Brains were fixed overnight at 4 °C in 4% PFA. Subsequently, tissues were cryo-protected using a sucrose gradient (30%) in PBS supplemented with 0.02% sodium azide (Sigma). Cryo-sectioning was performed using a cryo-stat (HN525 NX, Epredia, Germany) at a thickness of 30 µm (coronal). To investigate the impact of chronic cranial RT and RZ treatments on the neurogenic niche function, serial coronal brain sections (2 to 3 sections per brain, 8 to 10 brains per group) through the hippocampal formation were subjected to free-floating immunofluorescence staining using established protocols. Doublecortin (DCX) staining was conducted to label newly born, immature neurons, employing a rabbit anti-DCX primary antibody (1:200; Abcam) and a donkey anti-Rabbit Alexa Fluor 568 secondary antibody. DCX-positive cells were visualized in red using fluorescence microscopy. The BrdU-NeuN dual-immunofluorescence-stained tissues were permeabilized to recover the BrdU antigen and stained with mouse anti-BrdU (1:200) and rabbit anti-NeuN (1:500) primary antibodies. Fluorescence color facilitated using donkey anti-mouse Alexa Fluor 488 (1:200) and donkey anti-rabbit Alexa Fluor 568 (1:500) secondary antibodies. BrdU^+^ cells were observed in green, and NeuN^+^ neurons in red. Activated microglia were labeled by IBA1-CD68 dual immunofluorescence staining. Coronal tissues were permeabilized using 0.3% Tween-20 (Sigma) in PBS, followed by treatments with 3% hydrogen peroxide (Sigma) and 10% methanol (Sigma) on ice. Tissues were then blocked with 4% bovine serum albumin (Jackson ImmunoResearch) and 0.3% Tween-20 in PBS, followed by overnight incubation with primary antibodies (rabbit anti-IBA1, 1:500; and rat anti-mouse CD68, 1:500). Fluorescence was developed using goat anti-rat Alexa Fluor 647 (1:1000) and goat anti-rabbit Alexa Fluor 488 secondary (1:500) antibodies. IBA1^+^ microglia were observed in green, and CD68^+^ sub-cellular puncta in magenta. To assess synaptic marker, 30-μm-thick sections underwent immunostaining for synaptophysin. Sections were initially washed in PBS (pH 7.4), followed by a 30-minute blocking step in 4% (w/v) BSA and 0.1% Triton X-100 (TTX, Sigma). Subsequently, sections were incubated for 24 hours in a primary antibody solution containing 2% BSA, 0.1% TTX, and mouse anti-synaptophysin (1:1,000). Following primary antibody incubation, sections were treated for 1 hour with a solution of goat anti-mouse IgG labeled with Alexa Fluor 647 (1:1,000). For NeuN-cFos dual-immunofluorescence staining, 1:500 rabbit anti-cFos and 1:500 mouse anti-NeuN primary antibody solutions made in 3% NDS in Tris-A buffer were used. The secondary antibodies included 1:350 donkey anti-mouse AF 488, and 1:350 donkey anti-rabbit AF 568. After thorough rinsing in PBS, sections were sealed in a *Slow Fade* antifade Gold mounting medium (Life Technologies) for microscopic analysis. Detailed protocols are provided in the **Supplemental Information** section.

Single (DCX and synaptophysin), and dual (BrdU-NeuN and CD68-IBA1) immunofluorescence-stained sections were imaged using a laser-scanning confocal microscope (Nikon Eclipse AX) equipped with a 40× oil-immersion objective lens (1.0 NA) and NIS element AR module (v4.3, Nikon). High-resolution (1024 to 2048p) z stacks (1 µm thick) were acquired through the 25-30 µm thick brain sections. Unbiased deconvolution for the fluorescent z stacks and in silico volumetric quantification was carried out as previously described and detailed in the **Supplemental Information** section. An adaptive 3D blind deconvolution method (ClearView, Imaris v9.2, BitPlane, Inc.) was used to deconvolute and enhance the fluorescence signal resolution with respective fluorescent wavelengths. The deconvoluted images were analyzed using a 3D algorithm-based Imaris module (v9.2). BrdU, PSD-95, and synaptophysin, NeuN, IBA1, and CD68 were 3D modeled using the surface-rendering tool to create individual glial, synaptic, or neuronal cell volume. Using an unbiased, dedicated co-localization channel, the number of BrdU^+^ cells co-labeled with NeuN^+^ neurons was individually enumerated. Similarly, IBA1 and CD68 co-localization was determined for activated microglia. The number of DCX^+^ cells was quantified using the spot analysis tool. All Imaris-based *in silico* analyses were conducted using automated batch processing modules, with uniform criteria applied for all experimental groups by an experimenter blinded to the group IDs to avoid bias.

### ELISA for BDNF quantification

To assess the impact of RZ on hippocampal BDNF levels, mice subjected to RT ± RZ treatment were euthanized four weeks after the initiation of RZ treatment, and BDNF enzyme-linked immunosorbent assay (ELISA) was conducted following established procedures. Brains were promptly extracted from the skull (N=6-10 mice per group), and the hippocampus was micro-dissected from each cerebral hemisphere. The micro-dissected hippocampi were flash-frozen in cryovials by immersion in liquid nitrogen and stored at -80°C until assayed. Each hippocampus was weighed and transferred into 500 μl of ice-cold lysis buffer (NPER, Neuronal Protein Extraction Reagent, ThermoScientific) containing sodium orthovanadate (0.5 mM, Santa Cruz), phenyl-methylsulfonyl fluoride (PMSF, 1 mM, Santa Cruz), aprotinin (10 μg/ml, Santa Cruz), and leupeptin (1 μg/ml; Santa Cruz). Subsequently, tissues were sonicated individually, centrifuged at 4 °C, and the supernatants were collected and diluted at 1:5 or 1:10 with ice-cold Dulbecco’s PBS (Gibco). The supernatants were acidified to pH 2.6 and then neutralized to pH 7.6. BDNF levels were quantified using a commercially available ELISA kit (E-EL-M0203, Elabscience Biotechnology) and uncoated ELISA plates (Nunc MaxiSorp, Biolegend). Colorimetric measurements were conducted at a wavelength of 450 nm using a microplate reader (BioTek, SynergyMx).

### Spatial transcriptomics, MERFISH

We used a commercial spatial transcriptomics Multiplexed Error-Robust Fluorescence *in situ* Hybridization (MERFISH) platform (MERSCOPE, Vizgen, Cambridge, MA). MERFISH was performed according to Vizgen’s protocol as described previously [25]. Three brain samples, each from the RT + Veh group with the RT + RZ group, were utilized for the MERFISH experiments. Briefly, sagittal cryosections were collected from fresh-frozen OCT-embedded hemibrains of mice examined using a Leica CM1850 cryostat. The sections (10 um) were mounted onto Vizgen merslides (Vizgen #10500001) and hybridized with a specific panel of binary-coded probes that were custom-designed for 500 selected mouse genes (Vizgen #VZG0191, 40h at 37°C) after several steps of pretreatment including fixation (chilled 4% PFA in 1xPBS), permeabilization (70% ethanol), and prehybridization with formamide buffer (Vizgen). After hybridization, the brain sections on merslides were subject to washes with formamide buffer twice (30 minutes at 47°C), embedded with polyacrylamide gel, and cleared with a clearing mix containing protease K overnight at 37°C. Prior to MERFISH imaging, the cleared brain MERFISH samples were stained with DAPI-polyT staining kit included in the MERFISH imaging kit. The MERFISH imaging was conducted with a Vizgen MERSCOPE and a 500-gene imaging kit (Vizgen #10400006) after the merslide was uploaded. Imaging settings were consistent with those previously described[25] . Upon completion of imaging, the MERSCOPE program automatically converted the raw imaging data into *.vzg* meta output files, which were then used for downstream in-depth analysis using our customized bioinformatics pipelines.

### Statistical analysis

All data are expressed as the mean ± SEM. Statistical analyses of cognitive function, biochemical, and immunohistochemical data were conducted using two-way ANOVA (GraphPad Prism, v8.0). For the analysis of irradiation or RZ treatment effects, two-way ANOVA and Bonferroni’s multiple comparisons tests were applied. The exploration of familiar versus novel places in the OLM task by the same animals was compared using the Wilcoxon matched-pairs signed-rank test. Fear extinction training data were analyzed using repeated measures ANOVA and Bonferroni’s multiple comparisons tests. All statistical analyses were considered significant for a value of *P*≤0.05.

## RESULTS

### Riluzole treatment ameliorates cranial RT-induced cognitive dysfunction

Adult WT male mice received 9 Gy cranial RT and 48 h later administered RZ in drinking water (**Fig. 1A**). At 1-month post-RZ treatment, mice were handled, habituated, and tested on the cognitive function tasks (**Fig. 1B-F**). The open field activity of a cohort of animals in the open arena with bedding was monitored. We did not find significant differences in the percentage of time spent in the central zone (60% of the arena) for the spontaneous open field activity (**Suppl. Fig. S1A**). To determine if cranial RT and RZ treatment affected anxiety-like behavior, animals were administered an elevated plus maze (EPM) task that determines a preference for exploring either elevated open or closed arms of the maze under bright lights. We did not find significant overall group differences between groups for the percentage of time spent in the open arms (**Suppl. Fig. S1B**). These data indicate the absence of neophobic behavior during the spontaneous exploration cognitive testing. Animals were administered an object location memory (OLM) task to determine hippocampal function. During the OLM test phase, the comparison of the percentage of time animals explored the familiar and novel placements of objects (**Fig. 1C**) revealed significant differences for the Control + Vehicle, Control + RZ, and RT + RZ groups of mice (*P*<0.0001, 0.01 and 0.0001, respectively), but not for the RT + Vehicle mice, indicating that irradiated animals receiving vehicle were unable to differentiate between the familiar and novel locations. This behavior for the novel vs familiar location exploration is also reflected in the heat map of animal activity during the OLM test (**Fig. 1B**). This preference for the novel place exploration was then calculated as the Memory Index (MI). We found significant overall group differences between the experimental groups (**Fig. 1D**, F_(3, 66)_=6.43, *P*<0.001). Irradiated mice receiving Vehicle (RT + Vehicle) showed a significantly reduced preference to explore the novel location compared to the Control + Vehicle and RT + RZ groups (*P*<0.001 and 0.01, respectively). Importantly, irradiated mice receiving RZ treatment did not show a reduced MI, and exploration for the novel location was comparable to that of the Control + Vehicle and Control + RZ mice.

**Figure 1.**
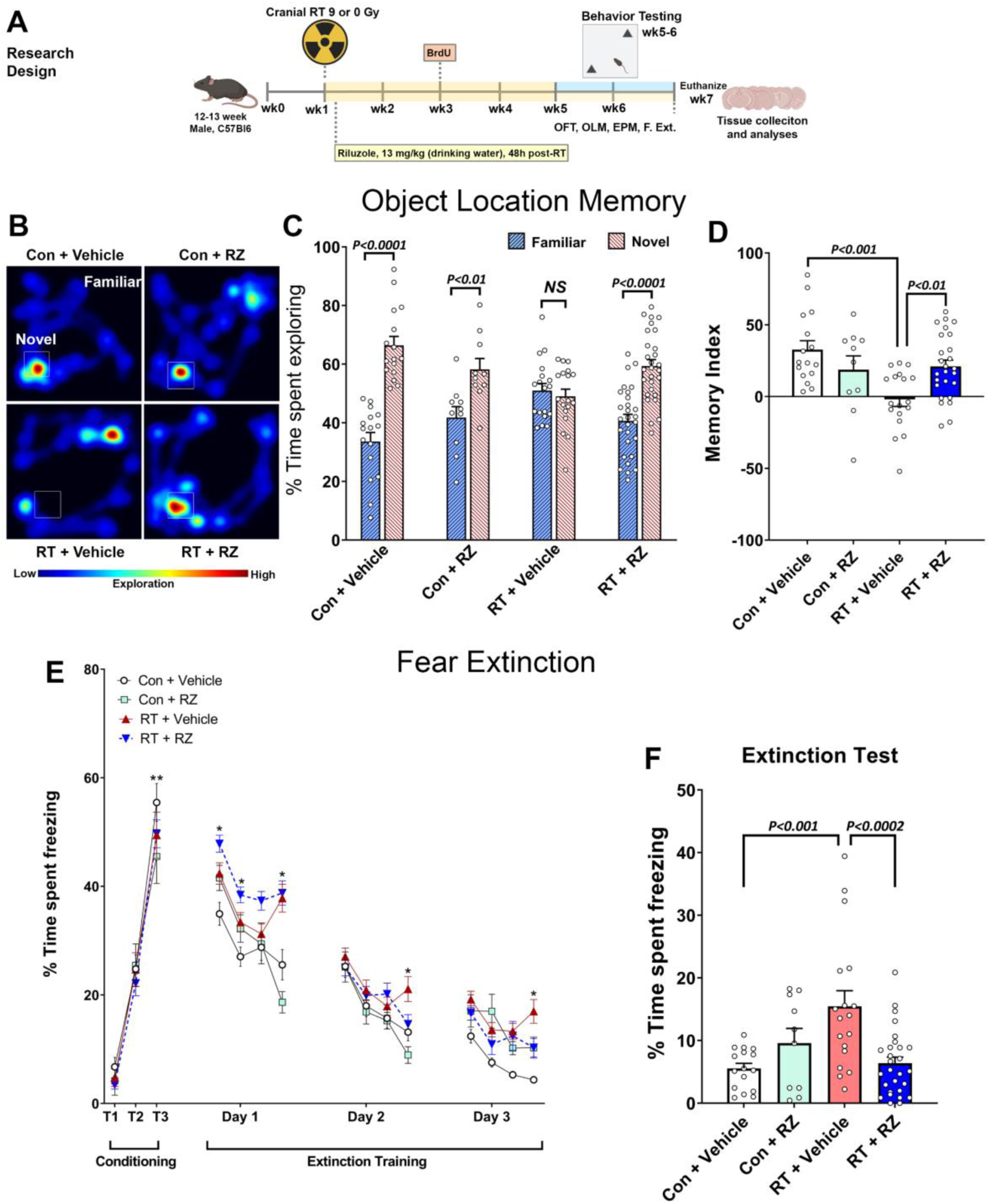
Treatment with riluzole reverses cranial radiation-induced cognitive impairments. **(A)** Research design: 12 to 13 weeks old wild type (C57BL/6) male mice received cranial radiation therapy (RT, 0 or 9 Gy) with protection of eyes and cerebellum. 48 hours after cranial RT, mice were treated with riluzole (RZ) in the drinking water (13 mg/kg) and continued on RZ till the end of the study (6-7 weeks). 0 Gy irradiated animals that received drinking water served as the vehicle group. For the assessment of dentate neurogenesis, two weeks post-RT, mice received BrdU injections (5-Bromo-2’-deoxyuridine, 50 mg/kg, IP, once daily for six days). One month after initiation of RZ treatment, mice were administered a hippocampal-dependent spatial memory retention test (Object Location Memory, OLM), anxiety-related tasks (Open Field Activity, OFT; and Elevated Plus Maze, EPM), and in the end, a fear extinction memory consolidation task (F. Ext.). After completion of cognitive testing, mice were euthanized and brains were collected for the tissue analyses. **(B)** Representative heat maps depicting animals exploring novel or familiar placement of objects in each experimental group during the OLM task. **(C)** Cranially irradiated mice receiving vehicle spent significantly less time exploring the novel placement of the object. Percentage time spent exploring the novel placements of objects during the test phase of the OLM task show that Control + Vehicle, Control + RZ and RT + RZ mice spent significantly more time exploring novel versus familiar location whereas RT + Vehicle group spent comparable time exploring both locations indicating a novel place location memory deficit. **(D)** For the OLM task, the tendency to explore a novel placement of the object was derived from the Memory Index (MI), calculated as ([Novel location exploration time/Total exploration time] – [Familiar location exploration time/Total exploration time]) × 100. Cranial RT significantly impaired spatial location memory as indicated by significantly reduced MI in the RT + Vehicle group compared to the Control + Vehicle, and RT + RZ group. Importantly, irradiated mice treated with RZ did not show a decline in spatial location memory and the DI was significantly higher compared to the RT + Vehicle group. **(E)** During the conditioning phase fear extinction memory task, cranial RT or RZ treatment did not impair the acquisition of conditioned fear response as indicated by the elevated freezing following a series of three-tone and shock pairings (80 dB, 0.6 mA, T1–T3). At 24 hours later, fear extinction training was administered every 24 h (20 tones) for the subsequent 3 days. Each data point for the extinction training Days 1-3 is presented as average of percentage time freezing for 5 tones (4 data points per day). All group of mice showed a gradual decrease in freezing behavior (Days 1–3), however, RT + Vehicle group spent a significantly higher time freezing compared with Control + Vehicle group. **(F)** Twenty-four hours after the extinction training phase, on the extinction test, Control + Vehicle and Control + RZ mice showed abolished fear memory (reduced freezing) compared with RT + Vehicle mice. Importantly, cranial RT-exposed mice receiving RZ (RT + RZ) were able to successfully abolish fear memory (reduced freezing) compared with the RT + Vehicle group. Data are presented as mean ± SEM (*N* = 10-26 mice per group). *P* values were derived from two-way ANOVA and Bonferroni’s multiple comparisons test. \**P* < 0.01, vs. Control + Vehicle. \*\**P* < 0.0001, T1 vs. T3 for all experimental groups.

Previously, we have shown cranial RT-induced deficits in contextual fear memory and fear extinction memory consolidation [35, 37]. To determine the beneficial neurocognitive impact of RZ, animals were administered the fear extinction memory consolidation task (FE, **Fig. 1E**). During the conditioning phase (Day 1), all treatment groups showed comparable associative learning as indicated by increased time spent freezing during the tone-shock conditioning phase (**Fig. 1E**; T_1_-T_3_; 45 to 50% on T_3_). Twenty-four hours after the conditioning phase, extinction training was administered (Days 1-3). Mice were presented with 20 tones per day (every 5 s intervals) in the same context and odor cue as the conditioning phase, but without foot shock. The RT + Vehicle group continued to show increased freezing compared to the Control + Vehicle group (**Fig. 1E**; Extinction Training Day 1-3, *P’s<*0.01). At 24 h after the completion of extinction training phase, animals were administered extinction testing (3 tones, 120s intervals, no foot shock) in the same testing environment as was used for extinction training (**Fig. 1F**). We found significant group effects in the extinction test (**Fig. 1F**; F_(3, 68)_=7.79, *P=*0.0022). RT + Vehicle mice failed to abolish fear memories during this retrieval testing phase as indicated by the increased freezing levels. On the contrary, RT + RZ mice showed reduced freezing levels compared to RT + Vehicle group (**Fig. 1F**; *P<0.0002*). These data indicated that RZ treatment to the cranially irradiated animals mitigated impairments in the ability to dissociate the learned response (freezing) to a prior aversive event (the tone-shock pairing). Overall, cognitive function testing demonstrated that RZ ameliorated cranial RT-induced cognitive deficits.

### Riluzole treatment augments hippocampal BDNF levels in the irradiated mice

The above cognitive function data indicate a neuroprotective impact of RZ treatment in the irradiated brain. Previously, we have shown RZ-mediated restoration of brain BDNF levels and reversal of cognitive impairments in a chemotherapy-related brain injury model [43]. RZ treatment was also beneficial in a mouse AD model [19]. To link improvements in cognitive function in the irradiated mice receiving RZ, we conducted an ELISA-based quantification of BDNF from the micro-dissected hippocampus. We found a significant overall group effect in the hippocampal BDNF levels (**Fig. 2**; F_(3, 26)_=8.61, *P<*0.001). Cranial RT (RT + Vehicle) was associated with 50-53% decrease in the BDNF levels compared with either Control + Vehicle (*P*<0.005) or Control + RZ (*P*<0.02) groups. RZ treatment significantly increased hippocampal BDNF levels in the irradiated mice (*P*<0.05) compared to RT + Vehicle group. Thus, we posit that RZ treatment augmented BDNF in the irradiated brain that potentially contribute to cognitive recovery.

**Figure 2.**
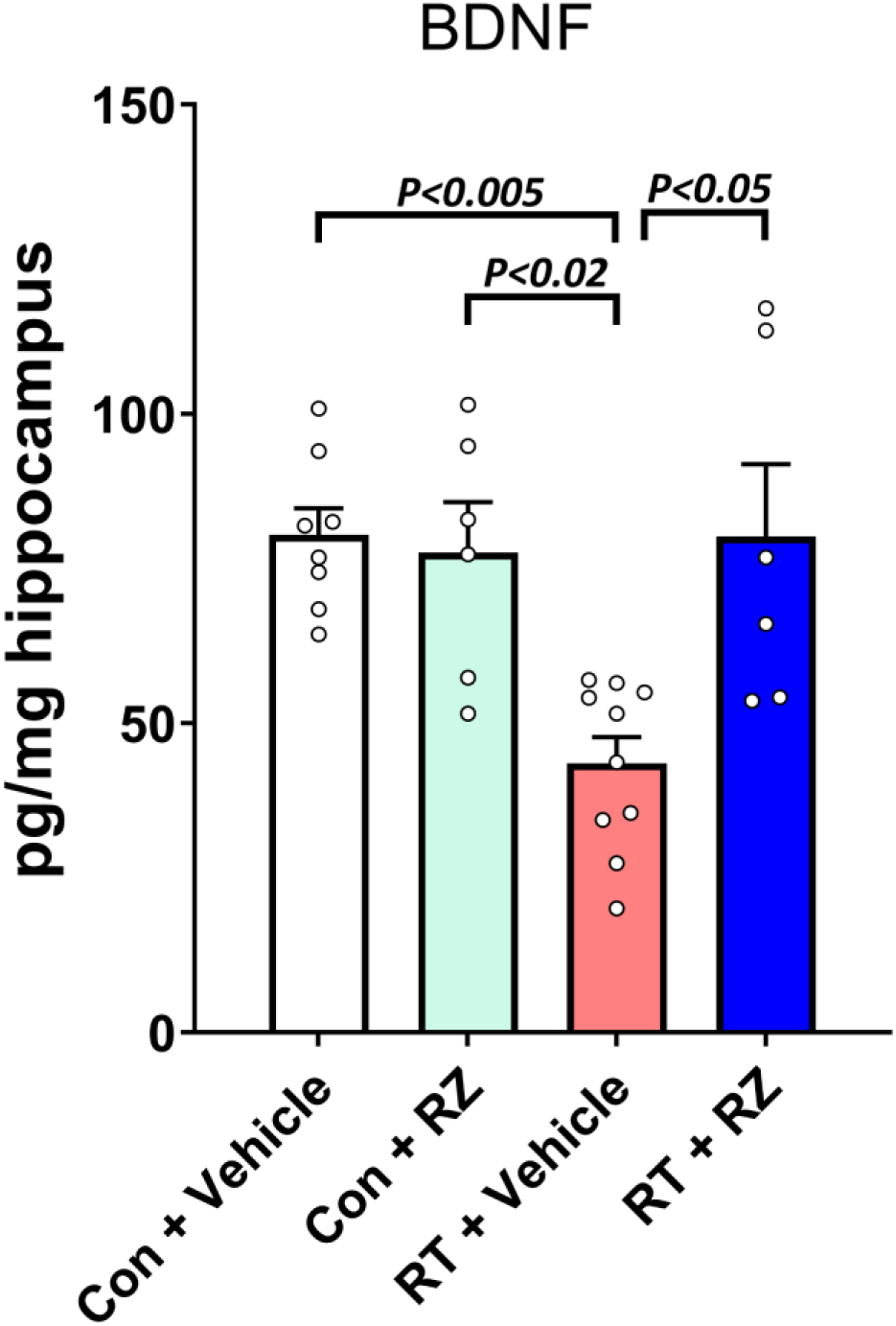
Riluzole treatment restores hippocampal BDNF in the irradiated hippocampus. 10-12 weeks old WT male mice received cranial radiation therapy (RT) followed by riluzole (RZ) treatment (13 mg/kg) in drinking water for 6-7 weeks. An ELISA-based quantification of BDNF from the micro-dissected hippocampus showed RT-induced reductions in the RT + Vehicle group. Importantly, RZ treatment in the cranially irradiated mice showed significant restoration of BDNF levels. Data are presented as mean ± SEM (*N* = 6-10 mice per group). *P* values were derived from two-way ANOVA and Bonferroni’s multiple comparisons test.

### BDNF enhancement restores cranial RT-induced synaptic loss

Our past work has shown cranial RT-induced loss of synaptic integrity including pre- and post-synaptic proteins that leads to long-term cognitive impairments [35, 37]. BDNF plays an important role in preserving neuronal function, including the synaptic landscape. To determine the impact of RZ treatment and BDNF augmentation on synaptic integrity in the irradiated brain, we quantified immunoreactivity of a synaptic marker, synaptophysin (**Fig. 3**). We found a significant overall group effect in the molecular layer (F_(3, 28)_=4.002, *P<*0.02) and stratum radiatum (F_(3, 28)_=19.32, *P<*0.0001) for the synaptophysin immunoreactivity. Cranial RT significantly reduced synaptophysin^+^ immunoreactive puncta compared to the Control + Vehicle group in the hippocampal dentate gyrus molecular layer (*P*<0.04, **Fig. 3A-B**) and CA1 stratum radiatum (*P*<0.0001, **Fig. 3C-D**). The synaptophysin levels were comparable between the Control + Vehicle and Control + RZ groups. Irradiated mice receiving RZ treatment did not show reduced synaptophysin compared to the RT + Vehicle group in the molecular layer (*P*<0.04) and stratum radiatum (*P*<0.001). Overall, these data indicate a protective role of RZ treatment and BDNF restoration against RT-induced disruption of synaptic loss.

**Figure 3.**
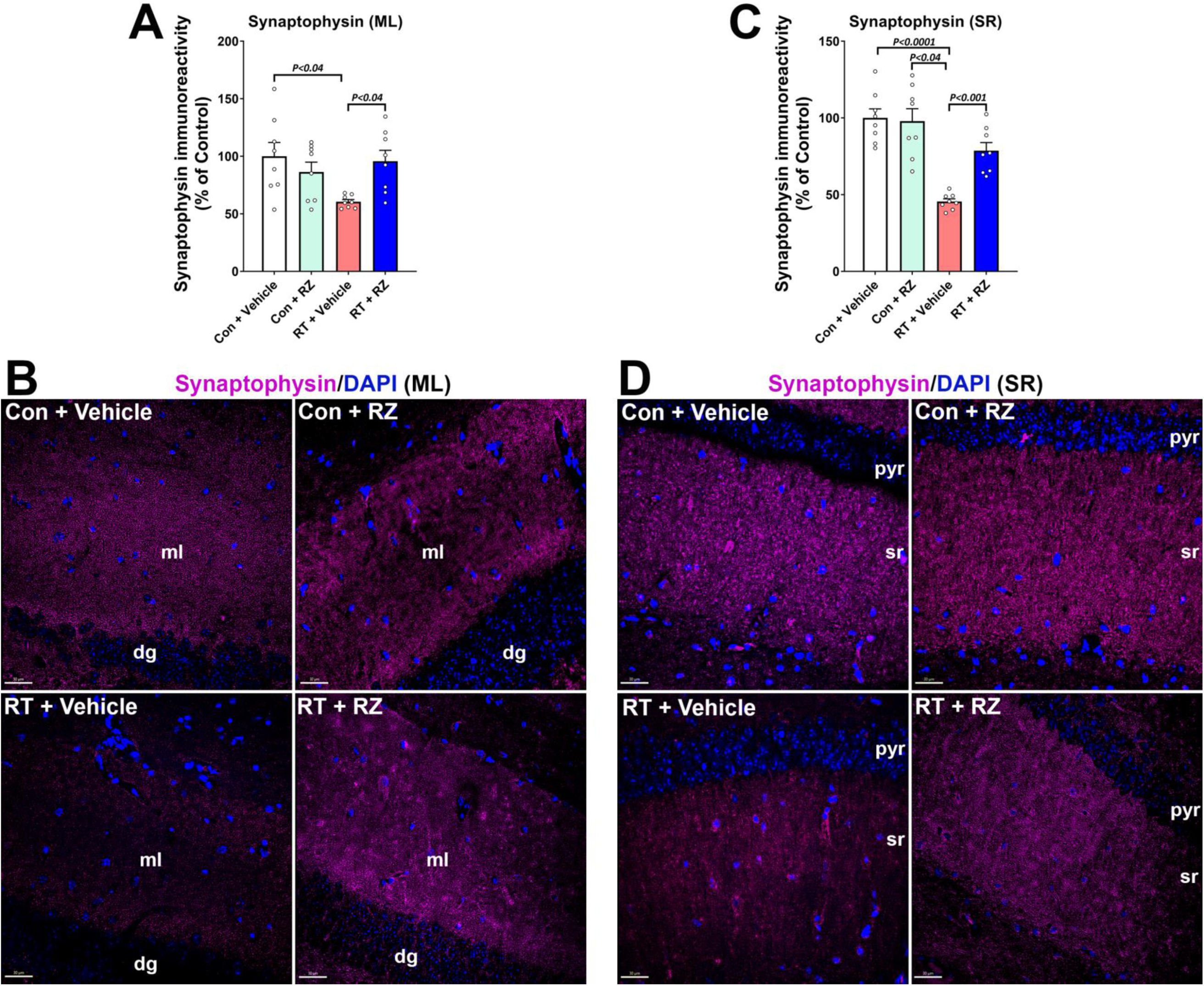
Riluzole treatment reverses cranial radiation-induced synaptic loss. Adult male mice were exposed to cranial RT and 48 h later received riluzole (RZ, 13 mg/kg) in drinking water for six to seven weeks. Synaptic protein marker, synaptophysin, in the molecular layer (ml) of hippocampal dentate gyrus (dg, **A-B**) and CA1 stratum radiatum (sr, **C-D**) emanating from the pyramidal neuron layer (pyr) was quantified using immunofluorescence staining, confocal microscopy and 3D algorithm-based unbiased fluorescent puncta analysis of synaptophysin immunoreactivity (magenta; DAPI nuclear stain, blue). Cranial RT (RT + Vehicle) significantly reduced the synaptophysin immunoreactivity in the dg ml, and CA1 sr regions compared with Control + Vehicle group. Irradiated mice receiving RZ showed a significantly increased synaptophysin immunoreactivity compared with the RT + Vehicle group. Data are presented as mean ± SEM (*N* = 8 mice per group). *P* values were derived from two-way ANOVA and Bonferroni’s multiple comparisons test. Scale bars, 30 μm, **B, D**.

### BDNF augmentation remediates cranial RT-induced reduction in neuronal plasticity

To further determine the impact RZ treatment and BDNF restoration in the irradiated brain on neuronal function, we evaluated the neuron plasticity-related immediate early gene (IEG) product, cFos, co-stained with the mature neuronal marker, NeuN, in the hippocampal dentate gyrus granule cell layer (GCL, **Fig. 4**). cFos facilitates neuronal LTP and hippocampal-dependent memory formation. A significant overall group effect was found in the number of cFos-NeuN^+^ dual-labeled cells (F_(3, 26)_=4.68, *P<*0.01). We found significantly reduced neuronal cFos (cFos-NeuN^+^) in the RT + Vehicle group compared to Control + Vehicle group (*P*<0.02, **Fig. 4A-D**) indicating compromised mature neuronal plasticity. Conversely, treatment with RZ significantly restored the number of cFos-NeuN^+^ cells in the irradiated GCL (**Fig. 4A, E**). This data in conjugation with synaptic protein expression data (**Fig. 3**) indicated restorative effects of RZ treatment in the irradiated brain.

**Figure 4.**
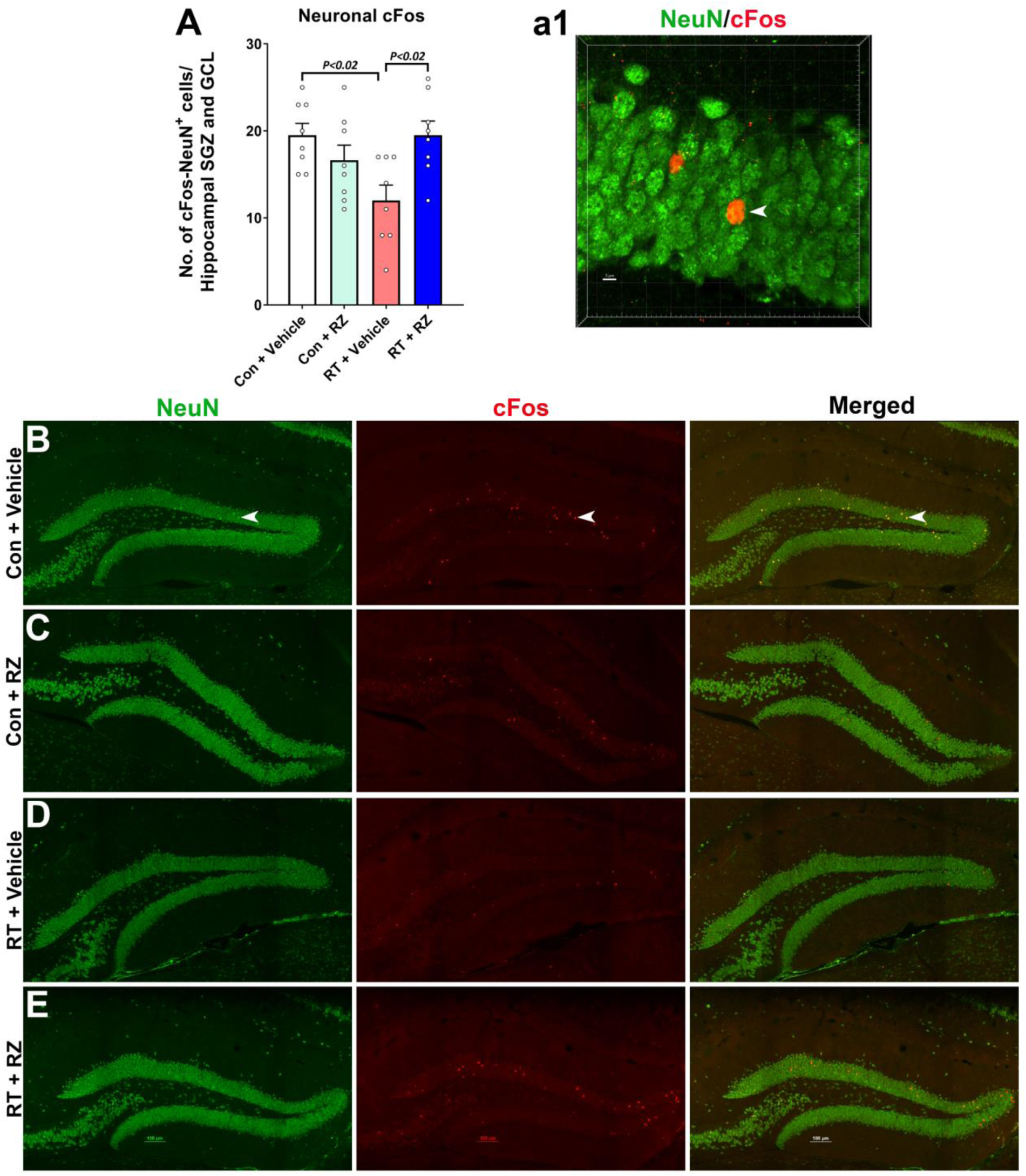
Treatment with riluzole prevents cranial radiation-induced reduction in the neuronal plasticity-related cFos expression. **(A, a1)** Dual immunofluorescence staining, confocal microscopy and 3D algorithm-based unbiased quantification of NeuN^+^ (green) neuronal plasticity-related immediate early gene (IEG) expression product, cFos showed a significant reduction in the number of cFos-NeuN^+^ neurons in the irradiated (RT + Vehicle) hippocampal sub-granular zone (SGZ) and the granule cell layer (GCL) compared to Control + Vehicle group. **(B-E)** Single channel fluorescent z stacks for NeuN (green), cFos (red) and merged images show cFos-NeuN co-expression in the hippocampal dentate gyrus for each treatment group. (**a1**) Higher magnification 3D fluorescent z stack of dual-stained neuron (arrow) is shown for the Control + Vehicle group (**B**). Treatment with riluzole significantly increased the number of cFos-NeuN^+^ neurons in the irradiated hippocampus (RT + RZ) compared to irradiated mice receiving vehicle (RT + Vehicle). Data are presented as mean ± SEM (*N* = 8 mice per group). *P* values were derived from two-way ANOVA and Bonferroni’s multiple comparisons test. Scale bars, 100 μm, **B-E**, and 5 μm, **a1**.

### Riluzole treatment did not reverse cranial RT-induced decline in neurogenesis

In our past study using a chemotherapy-induced cognitive impairment model, RZ treatment reversed chemotherapy-induced loss of newly born neurons and neurogenesis. Cranial RT has been shown to impact neural progenitor cell (NPC) function with an overall decline in doublecortin^+^ (DCX) newly born neurons and mature neurons differentiated from NPC pools in the hippocampal sub-granular zone (SGZ) [6, 8, 41]. We quantified DCX^+^ fluorescence neurons in the hippocampal GCL and SGZ. A significant overall group differences was found for the number of DCX^+^ immature neurons (**Fig. 5A-E**; F_(3, 24)_=255.5, *P<*0.0001). Cranial RT significantly reduced the number of DCX^+^ cells in hippocampal GCL and SGZ compared to either Control + Vehicle and Control + RZ groups (**Fig 5E**, *P*<0.001). Treatment with RZ did not mitigate the loss of DCX^+^ immature neurons in the irradiated hippocampus. Furthermore, we quantified the impact of RZ treatment on the dentate neurogenesis post-RT. Two weeks post-RT (**Fig. 1A**), mice were treated with BrdU to label proliferating NPCs. The assessment of neurogenesis (BrdU^+^-NeuN^+^ dual-labeled cells) was conducted 6-7 weeks later. We found overall significant effects on the percentage of BrdU^+^ cells expressing NeuN (**Fig. 5F-J**, F_(3, 51)_=70.22, *P<*0.0001). Cranial RT (RT + Vehicle) significantly reduced the percentage BrdU^+^-NeuN^+^ cells compared with Control + Vehicle and Control + RZ groups (**Fig. 5J**, *P*<0.001). Treatment with RZ did not restore BrdU^+^-NeuN^+^ cells in the RT + RZ hippocampus. These data indicate that in the irradiated microenvironment, treatment with RZ was ineffective in restoring neurogenesis.

**Figure 5.**
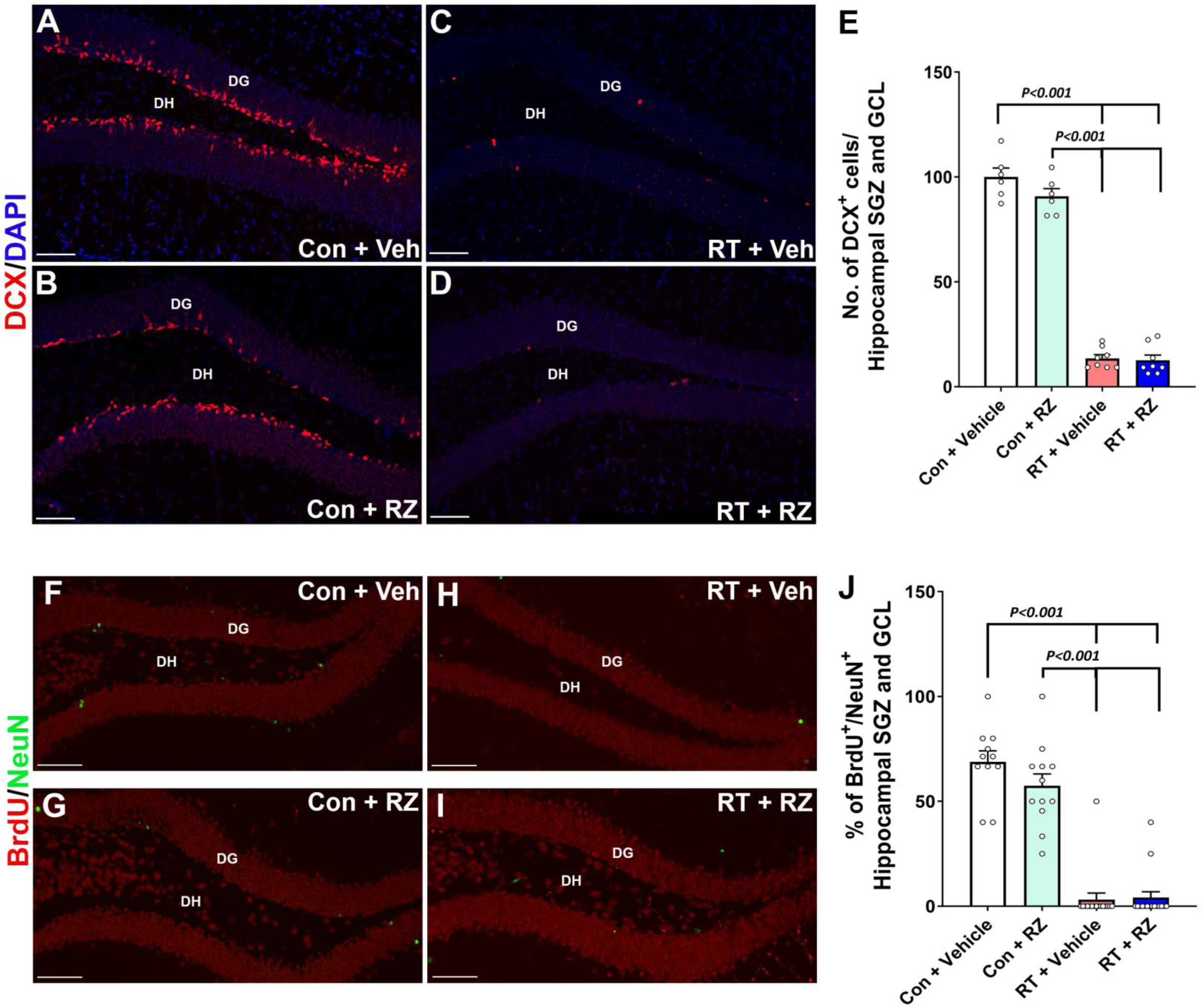
Riluzole treatment did not prevent cranial radiation-induced decline in neurogenesis. WT adult male mice received cranial RT (9 Gy) and treated with riluzole (RZ, 13 mg/kg) 48 h later in drinking water for six to seven weeks. Two weeks post-RT, mice were treated with BrdU and hippocampal neurogenesis was quantified using newly born neuron marker, doublecortin (DCX), and BrdU-NeuN dual-immunofluorescence staining in the hippocampal dentate gyrus (DG) sub-granular zone (SGZ) and molecular layer (ML) four to five weeks after the BrdU treatment. Cranial RT (RT + Vehicle) significantly reduced the number of DCX^+^ neurons (red, **A-D**) in the hippocampus compared with either Control + Vehicle or Control + RZ groups **(E)**. RT also significantly reduced neurogenesis, as indicated by the reduced percentage of BrdU^+^ cells (red) differentiating into the mature neurons (green, NeuN; **F-I**) in the RT + Vehicle group compared with either Control + Vehicle or Control + RZ groups (**J**). Riluzole treatment to the irradiated animals did not prevent the loss of DCX^+^ newly born neurons and the decline in dentate neurogenesis (BrdU^+^-NeuN^+^ dual-fluorescent cells). Data are presented as mean ± SEM (*N* = 6-16 mice per group). *P* values were derived from two-way ANOVA and Bonferroni’s multiple comparisons test. Scale bars, 50 μm, **A-D** and **F-I**.

### Riluzole treatment reduces cranial RT-induced neuroinflammation

Our previous studies, using chemotherapy and cranial RT rodent models [1, 2, 4, 5, 9, 35, 37], have shown elevated neuroinflammation, particularly microglial activation and astrogliosis, in the brain, linked with cognitive decline. In addition, we have shown that RZ treatment reduced microglial activation in the chemotherapy-exposed brain *in vivo* [5, 9, 13, 43]. Thus, to assess the effectiveness of RZ treatment on the status of microglial activation in the RT brain, dual immunofluorescence staining (**Fig. 6A-C**, and **a1**) and 3D algorithm-based volumetric analysis (**Fig. 6A**) were conducted for a pan microglial marker (IBA1) and a phagocytosis lysosomal marker (CD68). We found a significant overall group effect in microglial activation (IBA1-CD68 dual-immunoreactivity, F_(3, 24)_=12.95, *P<*0.001). Exposure to cranial irradiation significantly elevated IBA1-CD68 co-labeling in the hippocampus (RT + Vehicle, **Fig. 6A**) compared to Control + Vehicle (*P*<0.03) and Control + RZ (*P*<0.001) groups. Cranial RT also elevated astrocytic hypertrophy (astrogliosis) as indicated by thicker and longer stelae and GFAP immunoreactivity in the RT + Vehicle group compared to Control + Vehicle and Control + RZ groups (*P*<0.001, **Fig. 6D-E**). The overall group difference was significant for GFAP immunoreactivity (F_(3, 28)_=22.08, *P<*0.0001). On the other hand, RZ treatment to the irradiated mice (RT + RZ) significantly reduced activated microglia (IBA1-CD68; **Fig. 6A**) and astrogliosis (GFAP; **Fig. 6D**) compared to the RT + Vehicle group (*P<*0.001*, and P*<0.0001 respectively). These data signify the neuroprotective impact of RZ treatment in the irradiated brain leading to reductions in gliosis and cognitive dysfunction.

**Figure 6.**
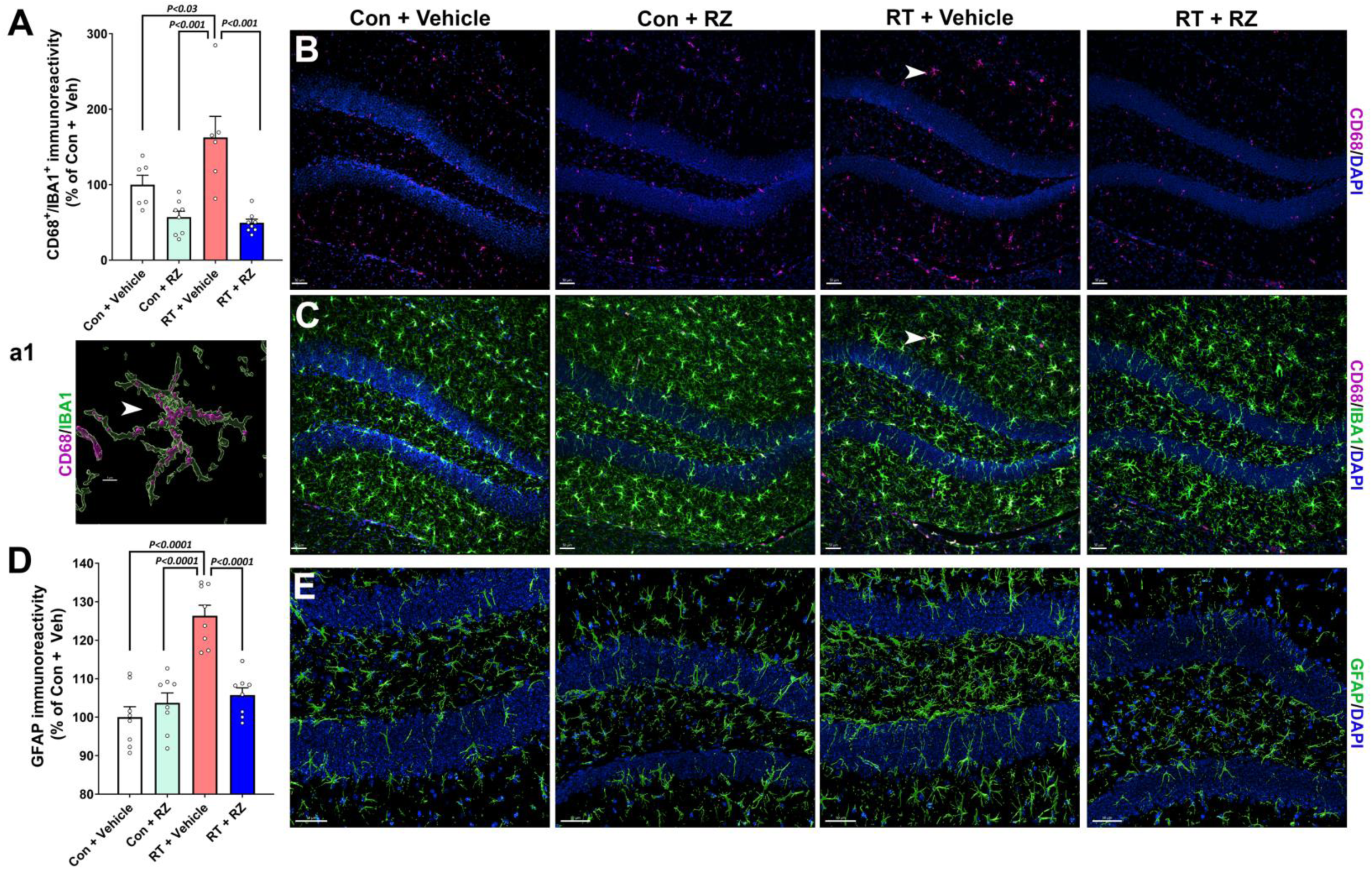
Riluzole treatment reduces cranial radiation-induced glial activation. Radiation-induced activated microglia in the hippocampal granule cell layer (GCL) and dentate hilus (DH) were assessed using dual immunofluorescence staining, laser scanning confocal microscopy and 3D algorithm-based volumetric quantification for IBA1 (green) and CD68 (magenta, and DAPI, blue) dual-labeling. **(A-C)** Exposure to cranial irradiation (RT + Vehicle) significantly elevated the immunoreactive volume of activated microglia (CD68^+^-IBA1^+^ co-localization) in the hippocampus compared with either Control + Vehicle or Control + RZ groups. Irradiated mice receiving RZ (RT + RZ group) showed a significantly reduced volume of CD68^+^-IBA1^+^ co-labeling compared with the RT + Vehicle group. CD68 alone **(B)** and CD68-IBA1 merged **(C)** fluorescence channels are shown for each group. **(a1)** Higher magnification surface reconstruction for the IBA1^+^ (green) positive microglia expressing CD68 puncta (magenta) is shown for the selected cell (white arrow, **B-C**) from the RT + Vehicle group. **(D-E)** Surface reconstruction of GFAP^+^ (green, DAPI, blue) hippocampal astrocytes shows significantly elevated GFAP immunoreactivity volume in the RT + Vehicle group compared to either Control + Vehicle or Control + RZ group. Astrocytes showed hypertrophy with thicker and longer stelae in the irradiated hippocampus (RT + Vehicle). RZ treatment to the irradiated mice (RT + RZ) showed a significant reduction in astrogliosis compared to RT + Vehicle group. Volumetric data for CD68^+^-IBA1^+^ co-expression and GFAP immunoreactivity are presented as mean ± SEM (*N* = 6-8 mice per group). *P* values were derived from two-way ANOVA and Bonferroni’s multiple comparisons test. Scale bars, 30 μm, **B**, 5 μm, **a1,** and 50 μm, **E**.

### Riluzole treatment improves transcriptomic signatures against cranial RT-induced neurodegeneration

For this study, we utilized MERFISH to perform spatial transcriptomics profiling of individual brain sections, with a specific focus on the hippocampal brain region [25, 49, 50]. Through the bioinformatics analysis platform, we collected about 6000 brain cells and identified 10 major cell types, including excitatory neurons from the CA1, CA3, and dentate gyrus regions, as well as glial cells including astrocytes and microglia within the hippocampus (Fig. 7A-B). Our dataset, which contains spatial locations and gene expression profiles of individual cells, including BDNF-related genes, combined with annotated cell types, offers a unique opportunity to explore the spatial *bdnf* and other gene expression in the intact tissue context. In the comparison of the RT + Veh with RT + RZ groups, we observed a significant increase in the *bdnf* gene expression in hippocampus excitatory neurons in the RZ-treated group (Fig. 7C-D, Wilcoxon Rank Sum test, average log2-fold change = 0.31, FDR-adjusted P<0.0001). Additionally, we found upregulated expression of an IEG, Nptx1, which is known to play a role in synaptic plasticity and neuroprotection, in hippocampus excitatory neurons in the RT + RZ group (Wilcoxon Rank Sum test, average log2-fold change = 0.27, FDR-adjusted *P*<10^16^). These findings suggest that RZ treatment counteracts cranial RT-induced neurodegeneration by enhancing neuroplasticity and neuroprotection in the hippocampus through the upregulation of key immediate early genes in excitatory neurons.

**Figure 7.**
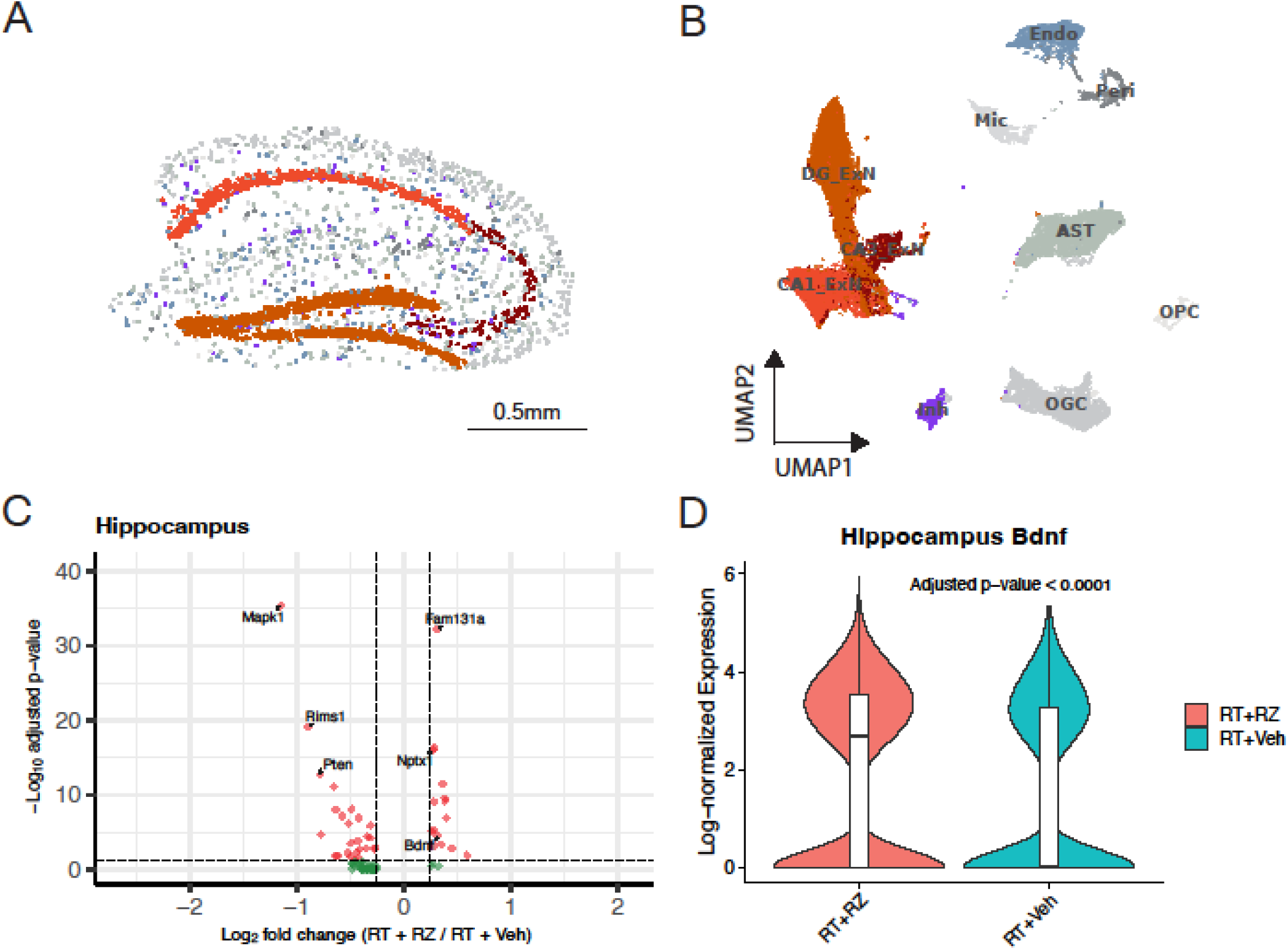
Spatial transcriptomic analysis of the cell-type-specific gene expression in the irradiated brain reveals upregulation of *Bdnf* and *Nptx1* in hippocampal excitatory neurons following riluzole treatment. **(A)** Spatial distribution plot of the hippocampus region in an individual brain section from the RT + Veh group, showing the localization of major cell types in MERFISH spatial transcriptomics profiling. **(B)** The UMAP visualization of 10 major cell types identified within the hippocampus, including excitatory neurons from the CA1, CA3, and dentate gyrus (DG) regions, inhibitory neurons, and non-neuronal (glial) cells including astrocytes (AST), microglia (Mic), oligodendrocyte granule cell and precursor cell (OGC, OPC), pericyte (Peri), endothelial cells (Endo). **(C)** Volcano plot illustrating differentially expressed genes between RT + RZ and RT + Veh groups in hippocampal excitatory neurons, highlighting the significant upregulation of Bdnf and Nptx1. **(D)** Violin plot showing the increased expression of Bdnf in hippocampal excitatory neurons of the RT + RZ group compared to the RT + Veh group. The expression values were log-normalized. P values were derived from Wilcoxon Rank Sum test and adjusted by Benjamini-Hochberg methods. Scale bar, 0.5 mm, **A**.

## DISCUSSION

This study verifies the neuroprotective role of riluzole treatment-mediated *in vivo* BDNF enhancement against cranial RT-induced cognitive dysfunction. Importantly, oral administration of RZ in the cranially irradiated mice lessened the RT-induced loss of synaptic integrity and neuronal plasticity and reduced neuroinflammation, hallmarks of radiation-induced long-term neurodegenerative consequences leading to cognitive decline. These results also corroborate our past findings showing the reversal of cognitive impairments and neurodegeneration in chemotherapy-induced brain injury [43]A likely common mechanism in each of these neurodegeneration and cognitive decline models was the reduction in BDNF. RZ treatment restored the neuronal BDNF *in vivo*, which played a contributory role in cognitive recovery in the irradiated brain.

Our hypothesis that enhancing levels of a neurotrophic factor in the irradiated brain would improve cognitive function is based on the scientific premise showing a correlation between BDNF levels and brain function. Behavioral deficits observed one month following a 30 Gy acute high-dose cranial RT was correlated with significantly reduced *bdnf* transcripts and protein expression [23]. These decrements in BDNF were linked with *bdnf* gene promoter histone H3 acetylation, a gene silencing epigenetic modification, in the irradiated hippocampus. Cranial RT doses of 5-10 Gy also downregulated BDNF mRNA, protein, and pCREB levels, phosphorylated cAMP-response element binding protein, in the irradiated hippocampus that was linked with hippocampal-dependent learning and memory deficits. Phosphorylation of CREB, a leucin-zipper class transcription factor, plays important roles prior to the transcription of neurotrophins (NTs) including BDNF and NT3 in hippocampal granule cell neurons [11]. Conversely, neuroprotective effects of BDNF-mediated cognitive recovery in a 20 Gy acute cranial RT model were blocked following treatment with a tyrosine kinase inhibitor that disrupts BDNF signaling [42]. These reports link cranial RT and the downregulation of BDNF expression and signaling impacting cognitive function. Interestingly, forced wheel running exercise following 20 Gy acute RT in rats improved cognitive function, neurogenesis and BDNF expression *in vivo* [22]. Physical exercise has also been shown to improve cognitive function following chemotherapy in animals [46]. Physical activities, including running, have been shown to be linked with increased brain BDNF levels that improve neuronal plasticity [44]. Collectively, these studies support an overall strategy for enhancing BDNF *in vivo* to improve cognitive function following cytotoxic cancer therapies. A number of studies also suggest a positive correlation between improved cognitive indices and BDNF in cancer patients undergoing chemotherapy [39], thus supporting our hypothesis that *in vivo* BDNF enhancement is neuroprotective against cranial RT-induced cognitive decline.

BDNF has been shown to play neuroprotective and functional roles in maintaining neuronal and dendritic health, activity-dependent neuronal plasticity, and cognitive function.[28, 33]. Reduced serum BDNF levels were associated with mild cognitive impairments, AD, and depression-related disorders [10]. Relatively higher expression of BDNF in the hippocampus has been linked with hippocampal-dependent cognitive function, synaptic plasticity and maintenance of neurogenesis [33]. Our past studies using cancer therapy-related cognitive impairment (CRCI) models demonstrated that exposure to either acute or fractionated RT and cytotoxic chemotherapy using drugs such as cyclophosphamide or doxorubicin significantly impaired dentate neurogenesis and cognitive function [6, 8, 13, 27, 30, 41]. Treatment with RZ in doxorubicin-exposed mice restored in DCX^+^ newly born neurons and neural progenitor cell (NPC)-derived mature neurons [43]. Conversely, in this study, RZ treatment and BDNF enhancement did not reverse RT-induced loss of neurogenesis (**Fig. 5**). Thus, in this study, RZ treatment-related improvements in cognitive function in the irradiated animals were unrelated to neurogenesis. To better understand the beneficial neurocognitive impact of RZ in the irradiated brain, we analyzed a pre-synaptic protein, synaptophysin, and a neuronal plasticity-related IEG product, cFos, expression in the irradiated hippocampus. Exposure to cranial RT significantly reduced synaptophysin in the dentate molecular and CA1 stratum radiatum layers **(Fig. 3)**. Our past studies using pre-clinical CRCI models have established that exposure to cranial RT, cytotoxic chemotherapy using cyclophosphamide or doxorubicin, and immune checkpoint inhibition therapy reduced hippocampal synaptic density that was associated with cognitive impairments [3, 5, 20, 27, 35, 37, 43]. Importantly, we also found significant reductions in IEG expression, including the neuronal cFos in the irradiated hippocampus **(Fig. 4)**. The cFos, similar to pCREB, facilitates hippocampal neuronal plasticity and LTP, glutamatergic receptor activity and memory function [26]. Conversely, treatment with RZ and *in vivo* increase in the hippocampal BDNF ameliorated RT-induced loss of synaptic integrity and neuronal IEG expression **(Fig. 4)**. Concurrently, using spatial transcriptomic profiling of hippocampus derived from the irradiated mice receiving RZ treatment, we found elevated Bdnf and Nptx1 gene expression within the excitatory neurons compared to irradiated mice receiving vehicle (RT + Veh, **Fig. 7**). NPTX1, neuronal pentraxin, is a crucial pre-synaptic class protein shown to facilitate synaptic strengthening, plasticity and resilience *in vivo* [14]. Loss of NPTX1 led to cognitive impairments in neurodegenerative conditions. Collectively, these data support the neuroprotective role of RZ treatment and BDNF augmentation in preserving mature, excitatory neuronal function in the irradiated brain and, thereby, cognitive recovery.

Our past pre-preclinical studies have shown cranial RT-induced microglial activation, astrogliosis, and elevated hippocampal cytokines (TNFa, IL1b, IL-6, IL-1a) that led to excessive synaptic loss and cognitive dysfunction [27, 35]. Our clinical studies of cancer patients undergoing chemotherapy found a correlation between pro-inflammatory signaling via TNFa, reduced plasma BDNF, and persistent cognitive impairments [40, 48], pointing to a neuro-modulatory role of inflammation and BDNF. Neurodegenerative condition-related neuroinflammation has been linked with the downregulation of the protective BDNF signaling [31]. In this study, we found reductions in the RT-induced hippocampal microglial activation and astrogliosis following RZ treatment and *in vivo* BDNF augmentation. Together, these data suggest that RZ treatment-related reductions in neuroinflammation in the irradiated brain contributed to preventing excessive synaptic and neuronal plasticity loss and cognitive decline.

While we have not evaluated these possibilities yet, we acknowledge that in addition to enhancing BDNF *in vivo* (as shown by ELISA, **Fig. 2**; and transcriptomics, **Fig. 7**), neuroprotective effects of RZ treatment can also positively impact other neural functions, including increasing LTP, reducing neuronal glutamate release, increasing glial glutamate reuptake, and inhibiting glutamatergic receptor activity that may influence irradiated brain function. Other reports demonstrating the beneficial effects of RZ on the AD brain have shown reduced neuronal glutamate release and amelioration of long-term memory deficits in AD mouse models. [16, 29] *In vitro* experiments using primary mouse astrocytes showed a 2-3 fold increase in BDNF, GDNF (glial cell line-derived neurotrophic factor), and NGF (nerve growth factor) synthesis 24 hours after initiation of RZ treatment [36]. Thus, both neurotrophic and neuronal mechanisms may contribute to the beneficial neurocognitive impact of RZ on the irradiated brain.

The overarching goal of our approach is to test this clinically-relevant, and translationally feasible, oral available, pharmacologic strategy in brain cancer-bearing mouse models receiving fractionated cranial RT and chemotherapy (*e.g.* temozolomide) regimes to expand the prospect of RZ treatment to ameliorate cognitive decline in cancer survivors and improve their quality of life. The future pre-clinical and clinical testing should also verify that treatment with RZ is safe in cancer-bearing subjects, and it does not interfere with therapeutic efficacy of cranial RT to thwart brain cancers. Nonetheless, this study provides proof of concept that *in vivo* augmentation of BDNF via oral administration of RZ in radiotherapy-injured brain is neuroprotective.

## CONCLUSION

Our data provides pre-clinical evidence for a translationally feasible approach: oral administration of an FDA-approved compound, riluzole, to enhance BDNF *in vivo* to ameliorate cranial radiation therapy-induced adverse impact on synaptic integrity, neuronal plasticity, neuroinflammation, and cognitive function.

## Supporting information

All_Supplemental_Figures

## ACKNOWLDEGMENTS

This work was supported by the National Institutes of Health (NIH) awards R01CA262213 (MMA) and R01CA276212 (MMA, AC), University of California Irvine (UCI) Institute of Clinical and Translational Sciences (ICTS) Pilot award through the NIH National Center for Advancing Translational Sciences (NCATS) award UL1TR001414 (MMA, AC). We also thank the support of the UCI CFCCC Biostatistics shared resource supported by the National Cancer Institute (NIH, award P30CA062203), and UCI Center for Neural Circuit and Mapping (CNCM). The content is solely the responsibility of the authors and does not necessarily represent the official views of the NIH.

## AUTHOR CONTRIBUTIONS

Conception and design: AC, MMA

Development of methodology: SMK, ARV, ACDL, ZT, XX

Acquisition of data: SMK, ARV, ACDL, ZT

Analysis and interpretation of data: SMK, DQN, JEB, ZT, XX, AC, MMA

Writing, review and/or revision of the manuscript: SMK, ARV, DQN, JEB, ZT, XX, AC, MMA

Administrative, technical, or material support: JEB, XX, AC, MMA.

Study supervision: JEB, XX, AC, MMA.

## DECLARATIONS

- Ethics approval: All animal experiments were approved by the UCI Institutional Animal Care and Use Committee (IACUC)
- Consent to participate: Not applicable
- Consent for publication: Not applicable
- Availability of data and material: All data and material are available per request
- Competing interests: The authors declare no competing interests

