## Supplementary material for "BDNF augmentation reverses cranial radiation therapy-induced cognitive decline and neurodegenerative consequences": All_Supplemental_Figures

**Supplemental Figure S1:**

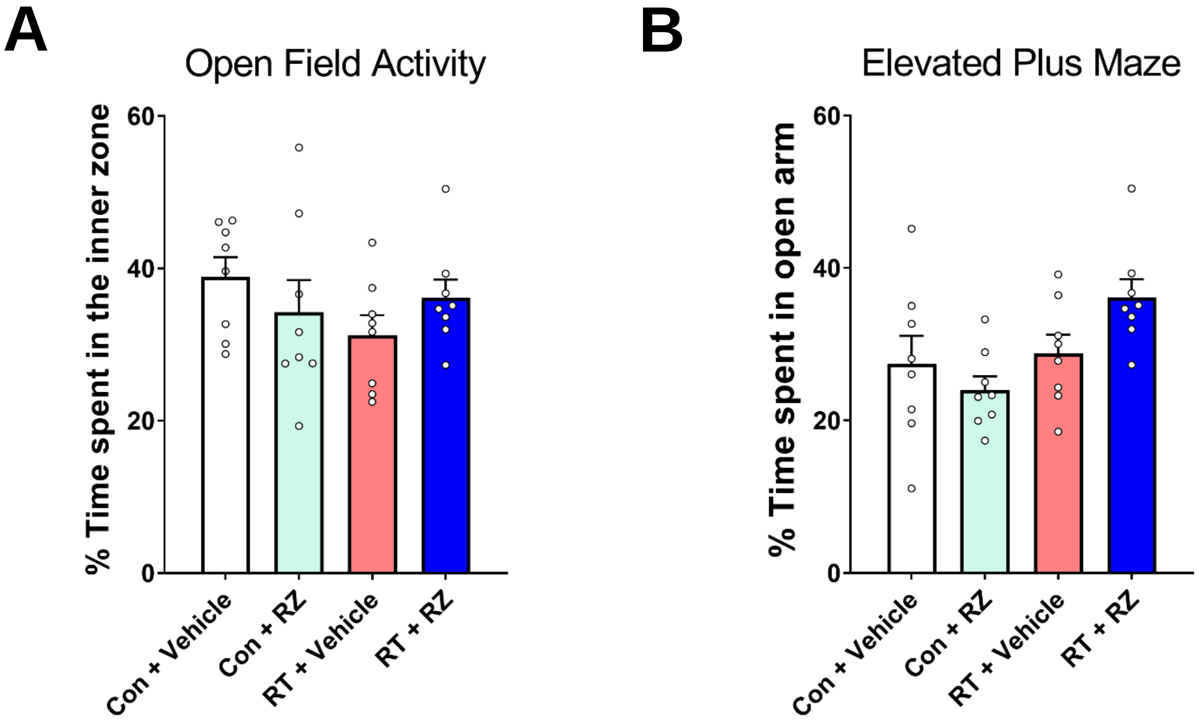


**Supplemental Figure S1:** Cranial radiation therapy (RT) or treatment with riluzole did not induce anxiety in animals. A cohort of mice from all groups were handled by the experimenter for one week prior to testing. The percentage of time spent in the central zone (**A**, 60% area) did not differ between the experimental group indicating the absence of neophobic behavior. For the elevated plus maze task, the percentage of time spent on the open arm (**B**) did not reach statistical significance between the experimental group indicating the absence of anxiety-related behavior. Data are presented as mean ± SEM (*N* =8 mice per group).

### RESOURCE AVAILABILITY

#### Lead Contact

#### Materials Availability

This study did not generate any new unique reagents.

#### Data and Code Availability

- All data reported in this paper will be shared by the lead contact upon request.
- This paper does not report the original code.
- Any additional information required to reanalyze the data reported in this paper is available from the lead contact upon request.

### EXPERIMENTAL MODEL AND SUBJECT DETAILS

#### Mice

All animals used in this study were cared for in accordance with the guidelines provided by NIH and approved by the Institutional Animal Care and Use Committee (IACUC) at the University of California, Irvine. Wild-type (WT) male mice, aged 12-13 weeks (C57BL/6J, RRID:IMSR_JAX:000664) were obtained from The Jackson Laboratory and housed in standard conditions (20 °C±1 °C; 70% ± 10% humidity;12h:12h light and dark cycle) in groups of 4 mice per cage. Littermates were randomly assigned to experimental groups, namely, Control-Vehicle (Veh), mice receiving 9 Gy cranial radiation therapy and vehicle (RT + Vehicle), 0 Gy irradiated mice receiving Riluzole (RZ) treatment (Con 9 RZ), and 9 Gy irradiated mice receiving RZ (RT + RZ).

### METHOD DETAILS

#### Experimental Design

Riluzole and its effects on cognition in the irradiated brain were studied in a mouse model. WT male mice were divided into the following groups: wild-type (WT) mice receiving 0 Gy or 9 Gy cranial radiation therapy (RT) and did or did not receive Riluzole (RZ) treatment. Treatment groups received RZ in drinking water 48 hours post-irradiation. BrdU injections were administered in all mice 2 weeks post-radiation for 6 days. After six weeks of continuous treatment with RZ, cognitive and behavior testing was performed, followed by euthanasia and harvesting of brains tissues for IHC, and ELISA.

**Cranial radiation therapy (RT)**

RT was administered to assigned experiment groups using the SmART+ image-guided research irradiator (Precision, Inc.). A total of 9 Gy head-only radiation was delivered to the anesthetized mice (5% induction and 2% maintenance isoflurane), with protection to eyes and cerebellum. RT was delivered through a 0.3 mm Copper filter, a 10x10 mm fixed collimator, and a photon spectrum of 225 kVp at 20 mA for 1 minute and 50 seconds at the region of interest.

**Riluzole preparation and administration**

Riluzole treatment was prepared as described previously (7). Riluzole (2-amino6-(trifluoromethoxy) benzothiazole, Selleckchem) was dissolved in filter sterilized (0.2 µm, Millipore), warm RO water (reverse osmosis, ULAR) with constant stirring (1 to 2 h at 45 °C) at a stock concentration of 600 µg per ml. The stock solution was kept frozen at −20 °C until used. The working solution (60 µg per ml) of RZ was prepared twice a week by diluting the stock with sterile RO water. Mice had ad libitum access to either vehicle (sterile RO water) or RZ solution (13 mg/kg per mouse) throughout the duration of the study.

**Open field activity**

To determine the spontaneous exploration of mice in an open arena, mice were allowed to explore arenas (33 × 33 × 33 cm) with bedding for 10 minutes. Open field activity was recorded as a percentage of time spent in the central (60% of central area) and the peripheral zones using Noldus EthoVision module (v17).

**Elevated Plus Maze (EPM) task**

The Elevated Plus Maze (EPM) is an anxiety assessment tool featuring a plus-shaped structure elevated above the ground. It includes two types of arms: one open with no walls or roof, and one closed with high walls, creating a darker, more enclosed environment. Before testing, the maze was cleaned with a deodorizer (Virkon, University Laboratory Animal Resources) and dried thoroughly. Mice were placed at the intersection of the arms and allowed to explore for 5 minutes, with their behavior recorded and analyzed using Noldus tracking software. Indices of percentage time spent in open arms served as an indicator of anxiety levels across different experimental groups.

**Object Location Memory (OLM) task**

Object location memory (OLM) task assesses spatial recognition memory function dependent on the integrity of the hippocampus (1). The experimental setup for OLM involved testing rooms with controlled lighting (50-70 Lux) and equipped with square arena boxes (33 × 33 × 33 cm), camera recording hardware (Noldus), and tracking software (EthoVision XT, v17). The tasks were conducted following previously established protocols (2,3). Mice underwent a habituation period where they were exposed to the open-field arena (with a thin bedding layer and no objects) for 10 minutes per day over three consecutive days. During the familiarization phase, mice explored two identical objects placed 16 cm apart for 5 minutes. After a 5-minute interval in the home cage, one object was moved to a new location within the arena, and the mice were allowed to explore for another 5 minutes. Behavioral data from all phases were recorded, and exploration time was analyzed using the "head direction to zone" function in EthoVision software. To ensure unbiased analysis, the time spent interacting with the familiar versus novel (or relocated) objects (with the mouse's nose within 2 cm of the object) was scored by observers blinded to the experimental conditions. The Memory Index (MI) for each animal was calculated using the equation: MI = ([Novel object or location exploration time/Total exploration time] – [Familiar object or location exploration time/Total exploration time]) × 100.

**Fear Extinction Memory**

To assess whether cranial RT or riluzole influences hippocampal circuit-dependent fear conditioning learning and fear memory consolidation processes, we conducted fear extinction behavior test (5,6). Testing occurred in a behavioral conditioning chamber (17.5 × 17.5 × 18 cm, Coulbourn Instruments) with steel shock floors (3.2 mm diameter slats, 8 mm spacing), and a waste collection tray sprayed with 10% vinegar. Mice were allowed to habituate in the chamber for 2 minutes during Day 1. Three pairings of an auditory conditioned stimulus (16 kHz tone, 80 dB, lasting 120 sec; CS) co-terminating with a foot shock unconditioned stimulus (0.6 mA, 1 sec; US) were presented at two-minute intervals. From days 2-4, all mice underwent initial 2-minute habituation followed by 20 non-US reinforced CS tones (16 kHz, 80 dB,lasting 120 sec, at 5 sec intervals). On the final day of fear testing (Day 5) mice were presented with only three non-US reinforced CS tones (16 kHz, 80 dB, lasting 120 sec) at two-minute intervals in the same spatial context and odor after the same two-minute habituation. Freezing behavior was recorded with a camera mounted above the chamber and scored by an automated measurement program (FreezeFrame, Coulbourn Instruments). FreezeFrame algorithms calculated a motion index for each frame of the video, with higher values representing greater motion. A blinded investigator established the motion index threshold for immobility for each animal individually by identifying a trough that distinguished low values (indicative of immobility) from higher values (associated with motion). Freezing behavior was defined as continuous immobility lasting one second or more. The percentage of time each mouse spent freezing was subsequently calculated for the final day of the extinction test.

**BDNF ELISA**

To assess the impact of RZ on hippocampal BDNF levels, mice subjected to RT and or RZ treatment were euthanized four weeks after the initiation of RZ treatment.  After which BDNF enzyme-linked immunosorbent assay (ELISA) was conducted following established procedures. Brains were promptly extracted from the skull (N = 6 to 10 mice per group), and the hippocampus was micro-dissected from each cerebral hemisphere. The dissected hippocampi were flash-frozen by immersion of the cryo-vials in liquid nitrogen and stored at -80°C until assayed. Each hippocampus was weighed and transferred into 500 μl of ice-cold lysis buffer (NPER, Neuronal Protein Extraction Reagent, ThermoScientific) containing sodium orthovanadate (0.5 mM, Santa Cruz), phenyl-methylsulfonyl fluoride (PMSF, 1 mM, Santa Cruz), aprotinin (10 μg/ml, Santa Cruz), and leupeptin (1 μg/ml; Santa Cruz). Subsequently, tissues were sonicated individually, centrifuged at 4 °C, and the supernatants were collected and diluted upto 1:10 ratio with ice-cold Dulbecco's PBS (Gibco). The supernatants were acidified to pH 2.6 and then neutralized to pH 7.6. BDNF levels were quantified using a commercially available ELISA kit (E-EL-M0203, Elabscience Biotechnology). Colorimetric measurements were obtained at a wavelength of 450 nm using a microplate reader (BioTek SynergyMx).

**Fixed Brains Collection**

Mice were anesthetized with isoflurane and euthanized via intracardiac perfusion using ice-cold saline (0.9% saline + 10 units/ml heparin) until the liver paled and venous outflow was clear. The saline-heparin buffer was then replaced with 4% paraformaldehyde (Sigma-Aldrich Cat. 158127, pH 7.4) and perfused until the body became rigid. Whole brains were extracted, soaked in 4% PFA overnight, and subsequently stored in PBS-0.05% sodium azide (Sigma-Aldrich Cat. S2002, pH 7.4). To prevent water crystallization during cryo-sectioning, brains were dehydrated in a sucrose gradient (10%, 20%, and 30% w/v, Sigma Cat. S7903, pH 7.4). Tissues were then embedded in O.C.T. compound (VWR Cat. 25608903) and cryo-sectioned (30 µm, coronal) using a Cryostat MX. Sections were stored in 24-well plates including PBS with 0.05% sodium azide (Sigma-Aldrich, pH 7.4) for future immunohistostaining.

**Immunohistochemistry**

Floating frozen brain sections were collected from each group (2-3 sections with mid-hippocampal region per brain, 4 brains per group) for immunohistochemistry. For synaptophysin, and cFos-NeuN tissue sections were washed in 1X TBS (dilute with MilliQ-Water from 10X TBS, Bioland Scientific Cat. TBS01-03, pH 7.4) three times (5 minutes each) and Tris-A solution (0.1% Triton-X, FisherSci Cat. 85111, in 1X TBS, ph 7.4) at room temperature (rt) for 10 minutes. Antigen retrieval for tissues from synaptophysin staining was facilitated by incubating in citrate buffer at 10 mM with 0.05% Tween-20 (pH 7.4 Sigma-Aldrich Cat. P4922 and Cat. 655204, respectively) at 70$^{o}$C for 30 minutes and recovering in borate buffer (100mM, Sigma, Cat. B0394, pH 8.5 ) at rt for 10 minutes and washed with 1X TBS (2 washes, 5 minutes each). Sections from were incubated in blocking solution with each respective host (3% normal goat serum, NGS) with 1% Bovine Serum Albumin or BSA in Tris-A buffer for Synaptophysin marker, and10% NDS in Tris-A for cFos-NeuN, NGS, Jackson ImmunoResearch Cat. 005-000-121; BSA, Sigma-Aldrich Cat. 05470) at rt for 45 minutes. Sections were incubated in primary antibodies at 4$^{o}$C overnight (1:1000 for Mouse Synaptophysin (Sigma Cat. S5768), in 3% NGS and 1% BSA in Tris-A buffer for Synaptophysin marker, and both 1:500 Rabbit anti cFos (Abcam Cat. ab190289), and 1:500 Mouse anti-NeuN, Millipore Cat. MAB377 with 3% NDS in Tris-A buffer for cFos NeuN marker). Before the second day of staining, sections were placed at room temperature on a shaker for 30 minutes. Sections were washed with 1X TBS (3 times, 5 minutes each) and then transferred to secondary antibodies to incubate at room temperature for one hour to visualize the target antigens (1:1000 Goat anti-Mouse AF 647, Abcam Cat. ab150115, for synaptophysin; 1:350 Donkey anti-Mouse AF 488, Invitrogen Cat. A21202, and 1:350 Donkey anti-Rabbit AF 568, Abcam Cat. ab175470 for cFos-NeuN marker). Lastly, brain sections were counterstained with DAPI nuclear dye (1µmol/L, FisherSci Cat. D1306) in 1X TBS at room temperature for 15 minutes. Stained sections were washed with 1X TBS (2 times, 5 minutes each) and mounted on superfrost slides (FisherSci Cat. 22-037-246) using VectaShield antifade mounting medium (VectaShield Cat. H-1000-10). Staining for the Doublecortin (DCX) marker followed similar steps as described above for cFos-NeuN with a minor change of buffer use (1X PBS, FisherSci), and antibodies (Primary antibodies: 1:750 Rabbit anti DCX, Abcam Cat. ab18723; Secondary antibodies: 1:500 Donkey anti Rabbit AF 568, Abcam Cat. ab175470).

Immunohistochemistry staining for CD68/IBA1 markers follow similar steps above with some minor changes. First, brain sections were washed with PBS-0.3% Tween 20 (Sigma-Aldrich) three times (5 minutes each) and then incubated in 3% peroxide solution (dilute 30% Hydrogen Peroxide, FisherSci Cat. H325-500, with 1% Methanol, FisherSci Cat. A412500), in 1X PBS, FisherSci) at room temperature for 30 minutes in order to block any endogenous peroxidase activity. Sections were washed in 1x PBS (3 times, 5 minutes each) prior to incubating in blocking solution (4% BSA in PBS-03% Tween-20, Sigma-Aldrich) at room temperature for 30 minutes. Brain sections were then incubated in primary antibodies at 4$^{o}$C overnight (1:500, Rat anti-Mouse CD68, Bio-rad Cat. MCA1957, combined with 1:500 rabbit anti-IBA-1, FUJIFILM Wako Cat. 019-1974, for CD68/IBA1 markers, with 1% BSA in PBS-0.3% Tween-20 buffer). Sections were washed and stained in secondary antibodies at room temperature for one hour (1:1000, goat anti-rat AF 647, Abcam Cat. ab150159, for CD68 marker, and 1:500, goat anti rabbit AF 488, FisherSci Cat. a11008, for IBA-1 marker). Sections were washed and counterstained with DAPI nuclear dye (1µmol/L, FisherSci Cat. D1306) and mounted on superfrost slides (FisherSci Cat. 22-037-246) using VectaShield antifade mounting medium (VectaShield Cat. H-1000-10).

Immunohistochemistry staining for BrdU-NeuN markers was performed similarly with the following changes. First, brain sections were washed with 1X PBS (FisherSci) buffer two times (5 minutes each) and then incubated in ice-cold 10% Methanol (FisherSci) for 20 minutes. Sections were then washed 3 times with PBS buffer (5 minutes each) followed by treatments with 50% formamide (ThermoSci) at 65 degrees C for 2 hours. Sections were washed three times with Saline-Sodium Citrate (SSC; Sigma-Aldrich) buffer (5 minutes each). Antigen retrieval for tissues was facilitated by incubating in borate buffer (100mM, Sigma, Cat. B0394, pH 8.5 ) at rt for 15 minutes and washing with 1X PBS (2 washes, 5 minutes each). Tissues were treated in block buffer (10% NDS in Tris-A) for 30 minutes and then placed in primary antibody (1:200 Mouse anti-BrdU, Millipore Cat. MAB3424, and 1:500 Rabbit anti-NeuN, Millipore Cat. ABN78) overnight. Steps were repeated as described above with the exception of the secondary antibody treatment (1:200 Donkey anti Mouse AF 488, Millipore Cat. MAB3424; 1:500 Donkey anti-Rabbit AF 568, Millipore Cat. ABN78).

**Confocal microscopy and 3D algorithm-based volumetric quantification**

Immunostained brain sections were captured at high resolution (1024p) using a laser-scanning confocal microscope (Nikon Eclipse AX) with a 40X oil-immersion objective lens, acquiring 0.5 µm thick z-stacks. The resulting high-resolution images were deconvoluted, and 3D volume surfaces for each antigen of interest were generated using Imaris (Andor Technologies), an advanced image analysis software. Synapse-related markers, such as synaptophysin, were quantified by analyzing the surface volumes of synaptic puncta. For cFos-NeuN, volumetric analysis was conducted using surface volumes of tagged cell bodies. For microglial markers like CD68/IBA1 and neurogenesis markers like, BrdU-NeuN, and cFos-NeuN, volumetric co-localization between the surfaces of the two markers was determined and analyzed. All image analyses were performed using automated batch processing, ensuring consistent application of parameters across all images for unbiased results.

| REAGENT or RESOURCES | SOURCE | IDENTIFIER |
|---|---|---|
| **Antibodies** | | |
| Rat anti M CD68 | BioRad | Cat. MCA1957; RRID: AB_322219 |
| Rabbit anti IBA1 | FUJIFILM Wako | Cat. 019-19741; RRID: AB_839504 |
| Mouse anti SV2 | DHSB | Cat. SV2 ; RRID: AB_2315387 |
| Mouse anti Synaptophysin | Sigma-Aldrich | Cat. S5768; RRID: AB_477523 |
| Mouse anti BrdU | Millipore | Cat. MAB3424 ; RRID:AB_94851 |
| Rabbit anti NeuN | Millipore | Cat. ABN78 ; RRID:AB_10807945 |
| Rabbit anti cFos | Abcam | Cat. ab190289 ; RRID:AB_2737414 |
| Mouse anti NeuN | Millipore | Cat. MAB377 ; RRID:AB_2298772 |
| Rb anti DCX | Abcam | Cat. ab18723 ; RRID:AB_732011 |
| Goat anti Rat AF 647 | Abcam | Cat. ab150159; RRID: AB_2566823 |
| Goat anti Rabbit AF 488 | FisherSci | Cat. a11008; RRID: AB_143165 |
| Goat anti Mouse AF 568 | Abcam | Cat. ab175473 ; RRID:AB_2895153 |
| Goat anti Mouse AF 647 | Abcam | Cat. ab150115; RRID: AB_2687948 |
| Donkey anti Mouse AF 568 | Invitrogen | Cat. A10037 ; RRID:AB_11180865 |
| Donkey anti Mouse AF 488 | Invitrogen | Cat. A21202 ; RRID:AB_141607 |
| Donkey anti-Rabbit AF 568 | Abcam | Cat. ab175470 ; RRID:AB_2783823 |
| **Chemicals, peptides, recombinant proteins, kits** | | |
| Riluzole | Selleckchem | Cat. S1614 |
| BrdU | Sigma-Aldrich | Cat. B5002 |
| Paraformaldehyde | Sigma-Aldrich | Cat. 158127 |
| Hydrogen Peroxide, 30% (Certified ACS) | FisherSci | Cat. H325-500 |
| Sodium azide | Sigma-Aldrich | Cat. S2002 |
| Sucrose | Sigma-Aldrich | Cat. S7903 |
| Tissue-Tek* O.C.T. Compound | VWR | Cat. 25608-930 |
| Tris-buffered Salin (10X TBS) | Bioland Scientific | Cat. TBS01-03 |
| Triton™ X-100 Surfact-Amps™ Detergent Solution | FisherSci | Cat. 85111 |
| Phosphate-Citrate Buffer with Sodium Perborate | Sigma-Aldrich | Cat. P4922 |
| TWEEN® 20 | Sigma-Aldrich | Cat. 655204 |
| Boric acid | Sigma-Aldrich | Cat. B0394 |
| Normal Goat Serum | Jackson ImmunoReasearch | Cat. 005-000-121 |
| Normal Donkey Serum | Jackson ImmunoReasearch | Cat. 017-000-121 |
| Bovine Serum Albumin | Sigma-Aldrich | Cat. 05470 |
| DAPI Nuclear Stain | FisherSci | Cat. D1306 |
| Methanol (Certified ACS), Fisher Chemical™ | FisherSci | Cat. A412500 |
| VectaShield antifade mounting medium | VectaShield | Cat. H-1000-10 |
| Formamide | ThermoFischer | Cat. 17899 |
| SSC buffer | Sigma-Aldrich | Cat. S6639 |
| Mouse BDNF Elisa Kit | Elabsciences | Cat. E-EL-M0203 |
